# Screening and identification of MicroRNAs expressed in perirenal adipose tissue during rabbit growth

**DOI:** 10.1101/737411

**Authors:** Guoze Wang, Guo Guo, Xueting Tian, Shenqiang Hu, Kun Du, Jingxin Mao, Xianbo Jia, Shiyi Chen, Jie Wang, Songjia Lai

**Affiliations:** Farm Animal Genetic Resources Exploration and Innovation Key Laboratory of Sichuan Province, Sichuan Agricultural University, Chengdu 611130, China; Guizhou Medical University, Guiyang 550025, China; College of Pharmacy and Biological Engineering, Chengdu University, Chengdu 610106, China; ChongQing University Of Science & Technology, Chongqing 401331, China

**Keywords:** Rabbit, MicroRNA, Adipose tissue, MiRNA-seq

## Abstract

MiRNAs regulate adipose tissue development, which are closely related to subcutaneous and intramuscular fat deposition and adipocyte differentiation. As an important economic and agricultural animal, rabbits have low adipose tissue deposition and are an ideal model to study adipose regulation. However, the miRNAs related to fat deposition during the growth and development of rabbits are poorly defined. In this study, miRNA-sequencing and bioinformatics analyses were used to profile the miRNAs in rabbit perirenal adipose tissue at 35, 85 and 120 days post-birth. Differentially expressed (DE) miRNAs between different stages were identified by DEseq in R. Target genes of DE miRNAs were predicted by TargetScan and miRanda. To explore the functions of identified miRNAs, Gene Ontology (GO) enrichment and Kyoto Encyclopedia of Genes and Genomes (KEGG) pathway analyses were performed. Approximately 1.6 GB of data was obtained by miRNA-seq. A total of 987 miRNAs (780 known and 207 newly predicted) and 174 DE miRNAs were identified. The miRNAs ranged from 18nt to 26nt. GO enrichment and KEGG pathway analyses revealed that the target genes of the DE miRNAs were mainly involved in zinc ion binding, regulation of cell growth, MAPK signaling pathway, and other adipose hypertrophy-related pathways. Six DE miRNAs were randomly selected and their expression profiles were validated by q-PCR. In summary, we provide the first report of the miRNA profiles of rabbit adipose tissue during different growth stages. Our data provide a theoretical reference for subsequent studies on rabbit genetics, breeding and the regulatory mechanisms of adipose development.

## Introduction

MicroRNAs (miRNAs) are endogenous non-coding RNAs, typically 18∼26 nucleotides in length, that regulate gene expression in eukaryotic cells. Mature miRNAs are produced from long primary transcripts through a series of nucleases that are further assembled into RNA-induced silencing complexes. These complexes recognize target mRNAs by complementary base pairing, leading to mRNA degradation and the inhibition of translation(Fabian et al. 2010). MiRNAs regulate a wide range of physiological processes, including growth and development, virus defense, cell proliferation, apoptosis and fat metabolism. Meanwhile, it has been well documented that MiRNAs regulate adipose tissue development, which are closely related to subcutaneous and intramuscular fat deposition(Guoxi et al. 2011; Guo et al. 2012) and adipocyte differentiation(Son et al. 2014).MiRNAs, including miR-27b(Karbiener et al. 2009), miR-103(Meihang et al. 2015) and miR-148a(Shi et al. 2015) regulate adipogenic processes, promoting or inhibiting adipogenesis in animals. This implicates miRNAs can be a new target for studying the molecular mechanisms governing fat development, growth and deposition in animals.

To-date, studies on the role of miRNAs during fat development have focused on humans, mice, livestock and poultry. Gu and colleagues (Gu and Eleswarapu 2007) screened miRNAs in bovine adipose tissue and breast tissue and identified 59 DE miRNAs, 5 of which differed from known mammalian miRNAs. Wang et al(Wang et al. 2018b) constructed an *in vitro* adipogenesis model of Crest-feather ducks and performed deep miRNA-sequencing, identifying 105 DE miRNAs, 12 of which were newly predicted and related to adipogenesis, including miR-223, miR-184-3p and miR-10b-5.

As an important economic and agricultural animal, rabbits are sources of meat and fur, and are widely used as experimental models in biomedical research. In addition, the adipose tissue of rabbits has low deposition rates during growth, making it an ideal model to study adipose regulation (Desando et al. 2013; Lunli et al. 2014; Wang et al. 2015; Ye et al. 2014; Yu et al. 2015). However, studies on the miRNAs related to fat deposition during the growth and development of rabbits are limited. In this study, we performed miRNA-sequencing during three important stages of fat deposition (35, 85 and 120 days post-birth), to identify key miRNAs that regulate adipose growth. Our findings provide a theoretical reference for subsequent studies on rabbit genetics and breeding and the regulatory mechanism of adipose development.

## Materials and methods

### Animal and sample collection

Tianfu Black rabbits (indigenous breed in Sichuan province of China) aged 35, 85 and 120 days were used in this study. Given the plasticity and maturation of rabbit adipose tissue, three biological replicates of perirenal adipose tissue were collected for 35 days (YR) and 120 days (TR), and two for 85 days (MR). The samples were snap frozen in liquid nitrogen, and stored at −80°C until RNA extraction.

### Total RNA extraction

Total RNA was isolated using Trizol Reagent (Life Technologies, Carlsbad, CA, USA). RNA purity and integrity were determined using a Nanodrop (Thermo Fisher Scientific, Waltham, MA, USA) and Agilent Bioanalyzer 2100 system (Agilent Technologies, CA, USA), respectively. Moreover, RNA concentrations were measured using a Qubit^@^ RNA Assay Kit and Qubit^@^ 2.0 Fluorometer (Life Technologies, Carlsbad, CA, USA). Only samples with RNA Integrity scores > 8 were used for sequencing.

### MiRNA library construction and sequencing

MiRNA libraries were constructed and sequenced by Mega Genomics Co.,Ltd., (Beijing, China). Sequencing libraries were prepared using TruSeq Small RNA Sample Prep Kits according to the manufacturers protocols (Illumina, San Diego, USA). Briefly, 3’ and 5’ linkers were used for cDNA synthesis, and PCR amplification. Target fragments were gel purified and the quality of the libraries were assessed using Bioanalyzer 2100 (Agilent, CA, USA). Libraries were sequenced on an Illumina Hiseq 2500 platform and 50-bp single-end reads were generated.

### MiRNA bioinformatics analysis

MiRNAs were analyzed using ACGT101-miR (LC Sciences, Houston, Texas, USA). The analysis procedure was as follows: (1) 3’ connector and non-specific sequences were removed to obtain clean data; (2) the length of the sequences were maintained at 18∼26nt through length screening; (3) mRNAs, RFam and Repbase databases were used for comparative analysis and the filtration of remaining sequences; (4) fitering was used to obtain effective data and precursors were compared to rabbit reference genomes (GCF_000003625.3_OryCun2.0_genomic.fa) for miRNA identification; (5) differentially expressed (DE) miRNAs were analyzed with p-value(FDR) ≤ 0.05 as the threshold; (6) target genes of DE miRNAs were predicted by TargetScan(Agarwal et al. 2015; Friedman et al. 2008; Nam et al. 2014) and miRanda(Betel et al. 2010; Doron et al. 2008); (7) GO functional annotation and KEGG Pathway analysis were used to investigate the functional enrichment of the identified miRNA target genes.

### Validation of DE miRNA by q-PCR

Primers for the miRNAs and internal controls (**Table 1**) were designed using Primer-BLAST (https://www.ncbi.nlm.nih.gov/tools/primer-blast/). MiRNA-specific primers were synthesized by Sangon Biotech Co. (Shanghai). Six DE miRNAs were reverse transcribed into cDNA using Mir-X™ miRNA First-Strand Synthesis Kits (Takara, Dalian, China) according to the manufacturer’s protocol. Q-PCR was performed using 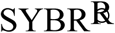 Green II qRT-PCR kits (Takara, Dalian, China) according to the manufacturer’s instructions. Reactions consisted of 4.5 μl 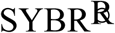 Green II, 1 μl cDNA, 0.5 μl of 10 μM forward and reverse primers, and 3.5 μl RNase free dH_2_O to a final volume of 10 μl. Reactions were performed on a Rotor gene 6000 PCR System (QIAGEN, Hiden, Germany) as follows: 95°C for 30 s, followed by 40 cycles of 95°C for 5 s, and 61°C for 20 s. The expression levels of the miRNAs were normalized to *GAPDH*. Relative gene expression was calculated using the 2^−ΔΔCt^ method(Livak and Schmittgen 2001). Data were expressed as the mean ± standard error of the mean (SEM).

**Table 1.**
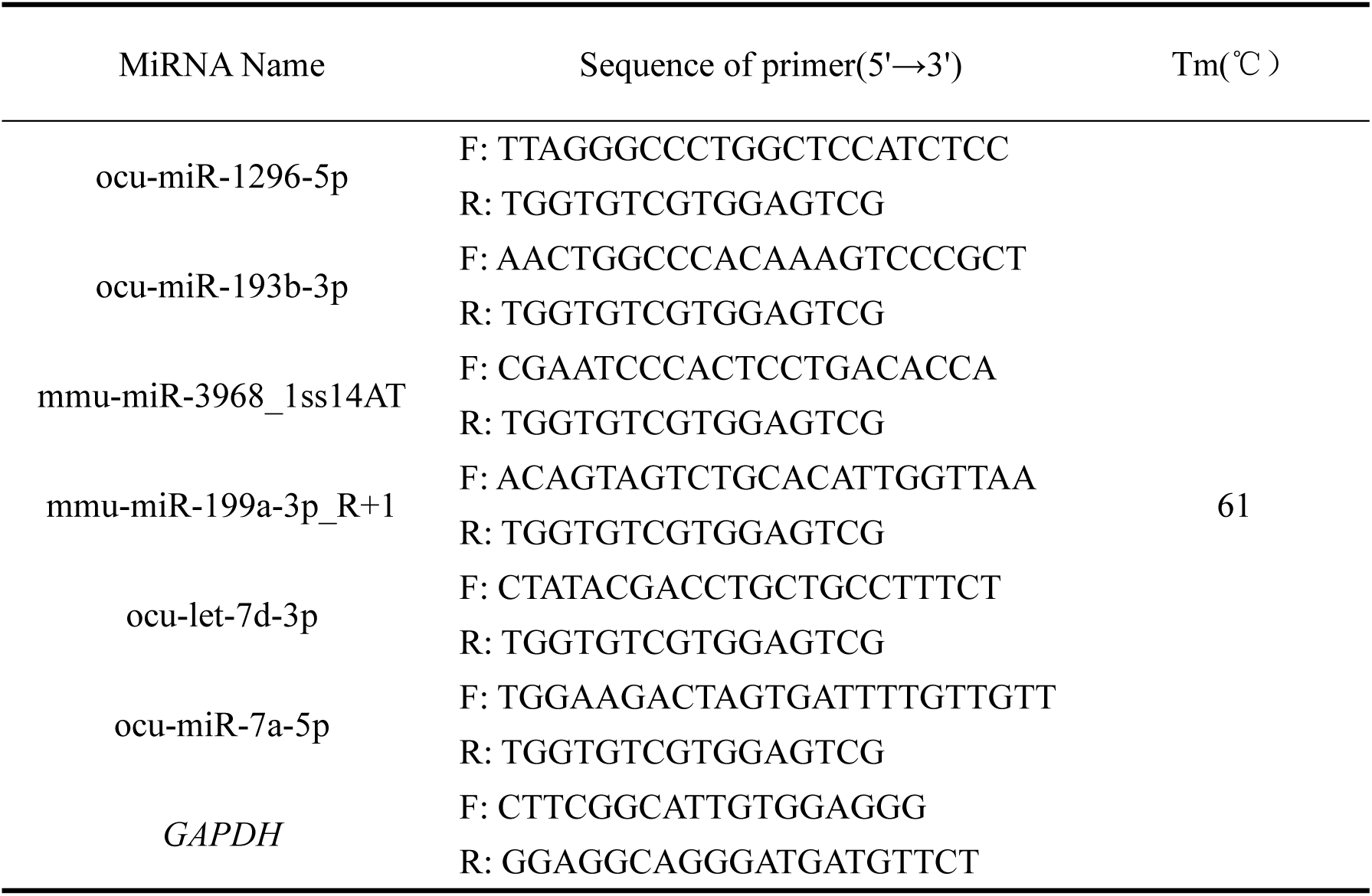
Primer information of 6 MiRNAs used for q-PCR validation.

### Statistical analysis

Statistical analysis was performed using SPSS Statistics 20.0 (SPSS Inc., Chicago, IL, USA). *P* < 0.05 was considered statistically significant.

## Results

### Overview of sequencing validation

Eight miRNA libraries of YR-1, YR-2, YR-3, MR-1, MR-2, TR-1, TR-2, TR-3 were constructed and divided into YR, MR, TR groups. Up to 1.6GB of data was obtained, and 8 libraries consisting of raw reads ranging from 10138426 to 15721988 were generated. FastQC (0.10.1) software was used to control data quality, through the removal of 3ADT & length filters (80% A or C / G or T; 3N; A alone; C without G; T alone; G alone; T without A; C alone; or continuous nucleotide dimers and trimers) and junk reads. After filtering and comparison to cellular mRNAs, RFam and Repbase databases, 1416639 ∼ 14139070 valid reads were obtained. The number of effective unique copies obtained from the libraries were 172905∼381169, accounting for 29.91%∼47.15% of the total sample (**Table 2**).

**Table 2.**
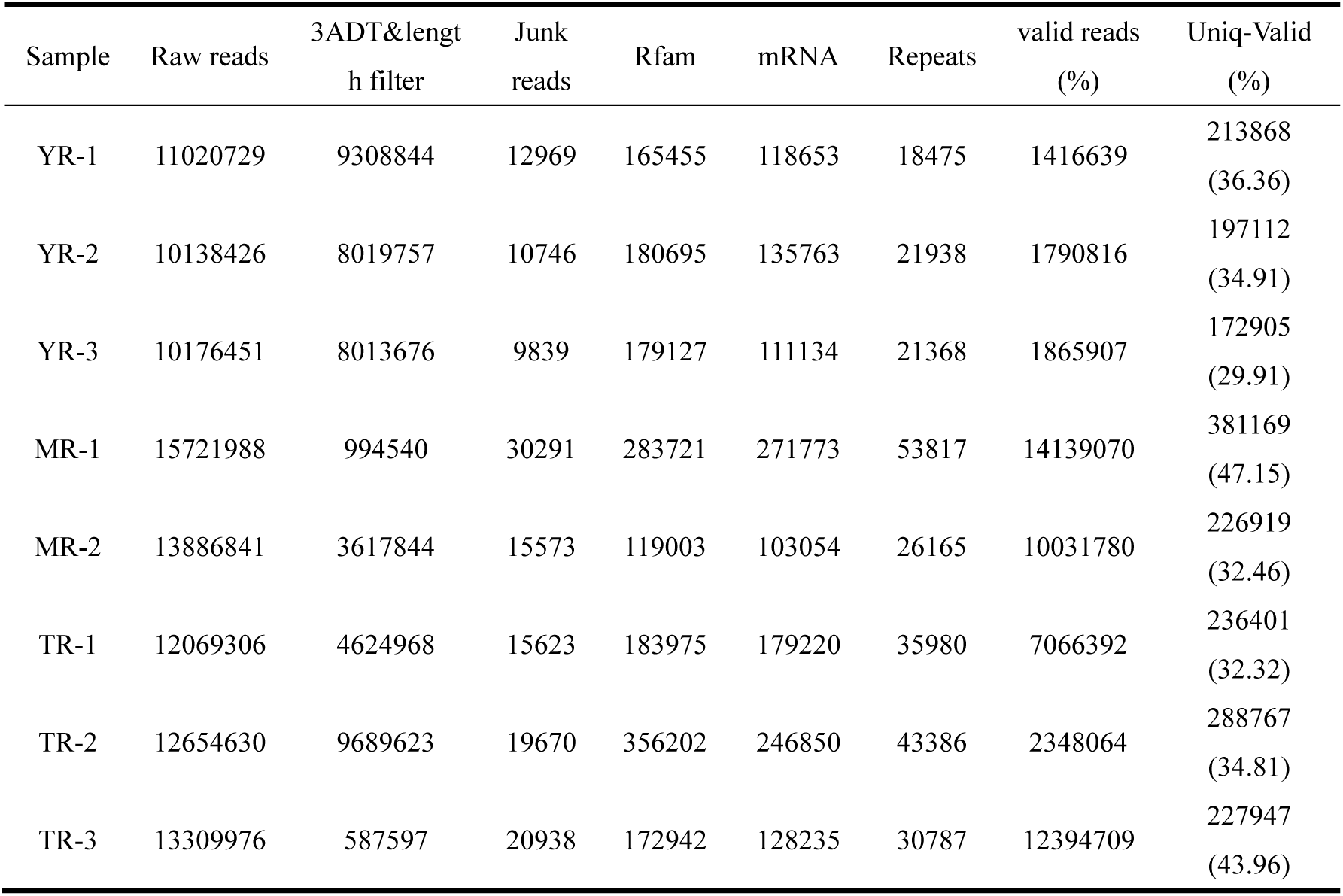
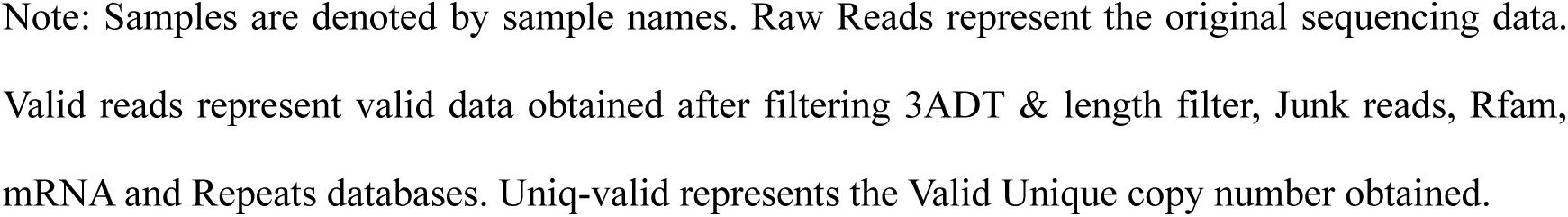
Summary and quality assessments of the sequencing data.

| Sample | Raw reads | 3ADT&length filter | Junk reads | Rfam | mRNA | Repeats | valid reads (%) | Uniq-Valid (%) |
| --- | --- | --- | --- | --- | --- | --- | --- | --- |
| YR-1 | 11020729 | 9308844 | 12969 | 165455 | 118653 | 18475 | 1416639 | 213868 (36.36) |
| YR-2 | 10138426 | 8019757 | 10746 | 180695 | 135763 | 21938 | 1790816 | 197112 (34.91) |
| YR-3 | 10176451 | 8013676 | 9839 | 179127 | 111134 | 21368 | 1865907 | 172905 (29.91) |
| MR-1 | 15721988 | 994540 | 30291 | 283721 | 271773 | 53817 | 14139070 | 381169 (47.15) |
| MR-2 | 13886841 | 3617844 | 15573 | 119003 | 103054 | 26165 | 10031780 | 226919 (32.46) |
| TR-1 | 12069306 | 4624968 | 15623 | 183975 | 179220 | 35980 | 7066392 | 236401 (32.32) |
| TR-2 | 12654630 | 9689623 | 19670 | 356202 | 246850 | 43386 | 2348064 | 288767 (34.81) |
| TR-3 | 13309976 | 587597 | 20938 | 172942 | 128235 | 30787 | 12394709 | 227947 (43.96) |

Note: Samples are denoted by sample names. Raw Reads represent the original sequencing data. Valid reads represent valid data obtained after filtering 3ADT & length filter, Junk reads, Rfam, mRNA and Repeats databases. Uniq-valid represents the Valid Unique copy number obtained.

### Length distribution of the candidate miRNAs

Following counting and analysis of the original sequencing data, the length distribution of the miRNAs in the 8 libraries were similar, varying from 18 nt∼26 nt, with 22nt miRNAs most frequent (**Figure 1**). To further analyze the validity of the sequencing data, statistics on the length distribution of miRNAs (Unique) were performed on filtered datasets. The results showed that number of the miRNAs in the 8 libraries were similar with > 60% of the reads 20∼24 nt in size, consistent with the characteristics of Dicer enzyme cleavage. Some miRNAs were in 25nt and 26nt in length, accounting for < 6% of the total sequences(**Table 3**).

**Table 3.** Sequence distribution of each unique MiRNA from each sample.

| length | YR-1 | YR-2 | YR-3 | MR-1 | MR-2 | TR-1 | TR-2 | TR-3 |
| --- | --- | --- | --- | --- | --- | --- | --- | --- |
| 18 | 28494<br>(13.32%) | 26300<br>(13.34%) | 26447<br>(15.30%) | 43900<br>(11.52%) | 17513<br>(7.72%) | 25208<br>(10.66%) | 43594<br>(15.10%) | 18368<br>(8.06%) |
| 19 | 28582<br>(13.36%) | 24865<br>(12.61%) | 24264<br>(14.03%) | 49220<br>(12.91%) | 18956<br>(8.35%) | 26278<br>(11.12%) | 42588<br>(14.75%) | 23592<br>(10.35%) |
| 20 | 27981<br>(13.08%) | 24640<br>(12.50%) | 22618<br>(13.08%) | 54436<br>(14.28%) | 21989<br>(9.69%) | 27892<br>(11.80%) | 40821<br>(14.14%) | 29560<br>(12.97%) |
| 21 | 30951<br>(14.47%) | 27716<br>(14.06%) | 26771<br>(15.48%) | 62982<br>(16.52%) | 33879<br>(14.93%) | 36155<br>(15.29%) | 44653<br>(15.46%) | 39681<br>(17.41%) |
| 22 | 26686<br>(12.48%) | 25794<br>(13.09%) | 22576<br>(13.06%) | 61580<br>(16.16%) | 36571<br>(16.12%) | 36073<br>(15.26%) | 37685<br>(13.05%) | 42513<br>(18.65%) |
| 23 | 24981<br>(11.68%) | 23443<br>(11.89%) | 15543<br>(8.99%) | 47535<br>(12.47%) | 30271<br>(13.34%) | 28478<br>(12.05%) | 27206<br>(9.42%) | 33886<br>(14.87%) |
| 24 | 34702<br>(16.23%) | 29210<br>(14.82%) | 27878<br>(16.12%) | 36777<br>(9.65%) | 49997<br>(22.03%) | 40183<br>(17.00%) | 38246<br>(13.24%) | 25985<br>(11.40%) |
| 25 | 7368<br>(3.45%) | 9123<br>(4.63%) | 4603<br>(2.66%) | 16040<br>(4.21%) | 11445<br>(5.04%) | 10603<br>(4.49%) | 9194<br>(3.18%) | 9908<br>(4.35%) |
| 26 | 4123<br>(1.93%) | 6021<br>(3.05%) | 2205<br>(1.28%) | 8699<br>(2.28%) | 6298<br>(2.78%) | 5531<br>(2.34%) | 4780<br>(1.66%) | 4454<br>(1.95%) |

Note: Length represents the length of MiRNA sequences.

**Figure 1.**
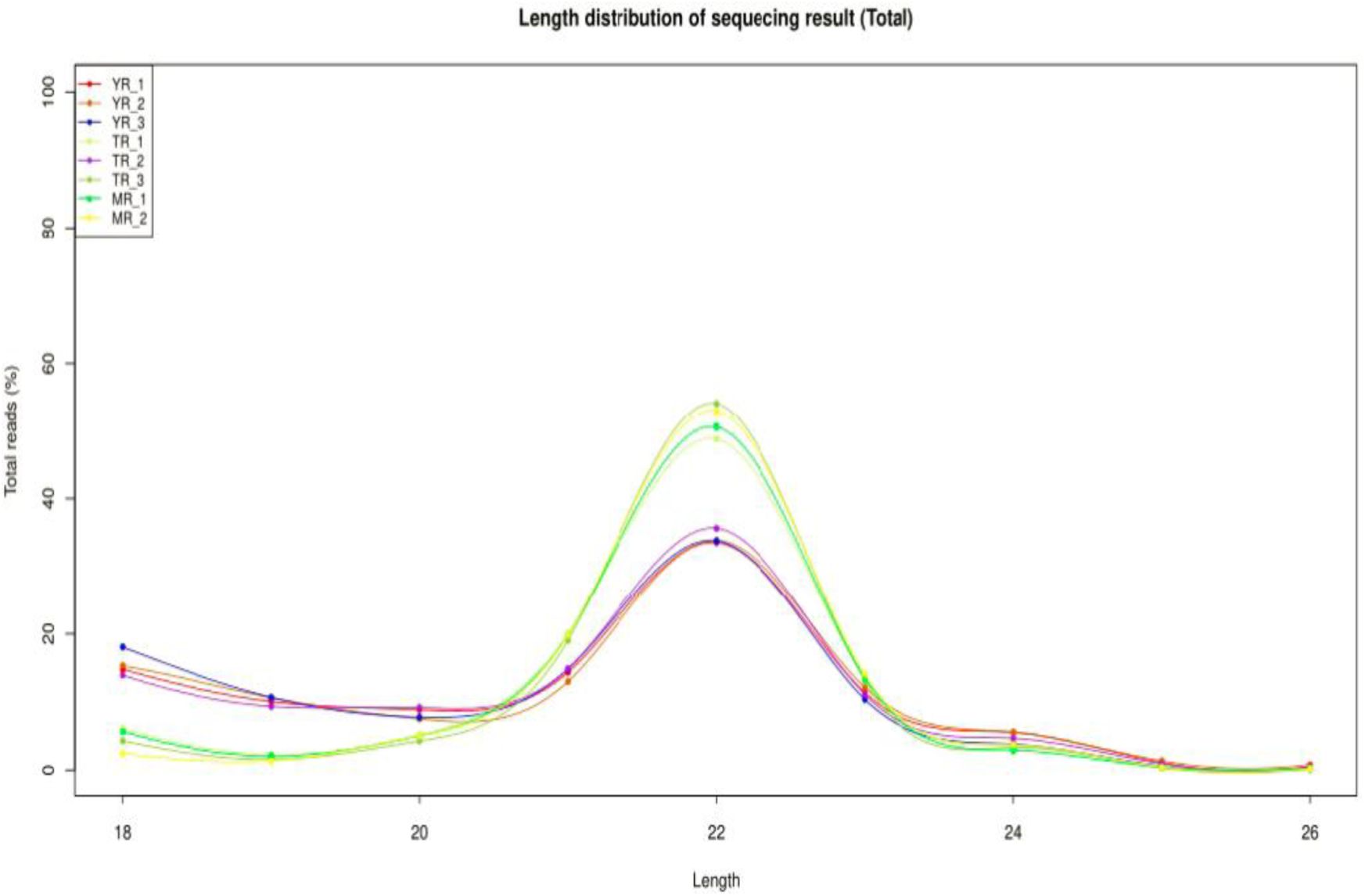
Length distribution of the 987 miRNAs.

### Annotation and identification of miRNAs

To obtain conserved miRNAs in rabbit adipose tissue, the ACGT101-miR (4.2) tool was used to compare the reference genome-matched reads with the known mature miRNAs in the miRase database. As a result, a total of 987 miRNAs were obtained during the three adipose growth stages, including 780 known miRNAs and 207 newly predicted miRNAs. Meanwhile, 131 miRNAs were highly expressed, 652 were moderately expressed, and 204 were expressed to low-levels. In addition, miRNA expression varied during different adipose growth stages, with 620 miRNAs obtained by YR (35 days), 865 obtained by MR (85 days), and 879 obtained by TR (120 days). These results showed that miRNA expression gradually increases during the adipose growth of rabbits.

As the sequence lengths of the miRNAs influence their regulation, the length distribution of the 987 miRNAs were assessed. The results showed that the lengths ranged from 18-26 nt, with 398 miRNAs 22 nt in length, accounting for the highest proportion (40.32%). while 26 nt miRNAs were least common (0.61%). The length distribution of the 780 known miRNAs was consistent with the total miRNAs, with the majority 22 nt in length (43.08%). of the 207 newly predicted miRNAs, none were 26nt, 2 were 25 nt, and 62 were 22 nt in length (**Table 4**).

**Table 4.** Length distribution of the identified MiRNAs.

| Length | Unique miRNA | Known miRNA | New miRNA |
| --- | --- | --- | --- |
| 18 | 40 (4.05%) | 20(2.56%) | 20(9.66%) |
| 19 | 37 (3.75%) | 8(1.03%) | 29(14.01%) |
| 20 | 44 (4.46%) | 27(3.46%) | 17(8.21%) |
| 21 | 180 (18.24%) | 154(19.74%) | 26(12.56%) |
| 22 | 398 (40.32%) | 336(43.08%) | 62(29.95%) |
| 23 | 212 (21.48%) | 170(21.79%) | 42(20.29%) |
| 24 | 55 (5.57%) | 46(5.89%) | 9(4.35%) |
| 25 | 15(1.52%) | 13(1.67%) | 2(0.96%) |
| 26 | 6(0.61%) | 6(0.76%) | 0 |
| all | 987 | 780 | 207 |

The 987 miRNAs were next analyzed to assess their evolutionary conservation. The results showed that miRNAs originated from 103 families, the numbers of which were differentially distributed. Members of the let-7 and miR-10 families were most frequent (11 miRNAs). Single miRNAs were identified for miR-196, miR-130 and miR-205 families.

### Identification of differentially expressed miRNAs

The DEGseq package in R was used to identify DE miRNAs and adjusted *P*-values (FDR) ≤ 0.05 were taken as standards to screen DE miRNAs during the three stages of rabbit adipose growth. A total of 174 DE miRNAs were obtained from 987 miRNAs in the three groups, of which 40.4% were up-regulated and 59.6% were down-regulated, indicating that the proportion of down-regulated miRNAs during rabbit adipose growth was significantly higher than the number of up-regulated miRNAs. Pairwise comparisons of the YR, MR and TR miRNA data showed 7, 164, 12 DE miRNAs between the respective growth stages, amongst which the number of DE miRNAs in the YR-vs-MR comparison group were the largest (**Figure 2**). Through in-depth analysis of the miRNA data obtained from inter-group comparisons, 12 DE miRNAs of YR-vs-TR showed moderate expression, 3 DE miRNAs of TR-vs-MR showed moderate expression, and 49 DE miRNAs showed high expression in YR-vs-MR, indicating that miRNA expression was more active at 85 day of rabbit adipose growth.

**Figure 2.**
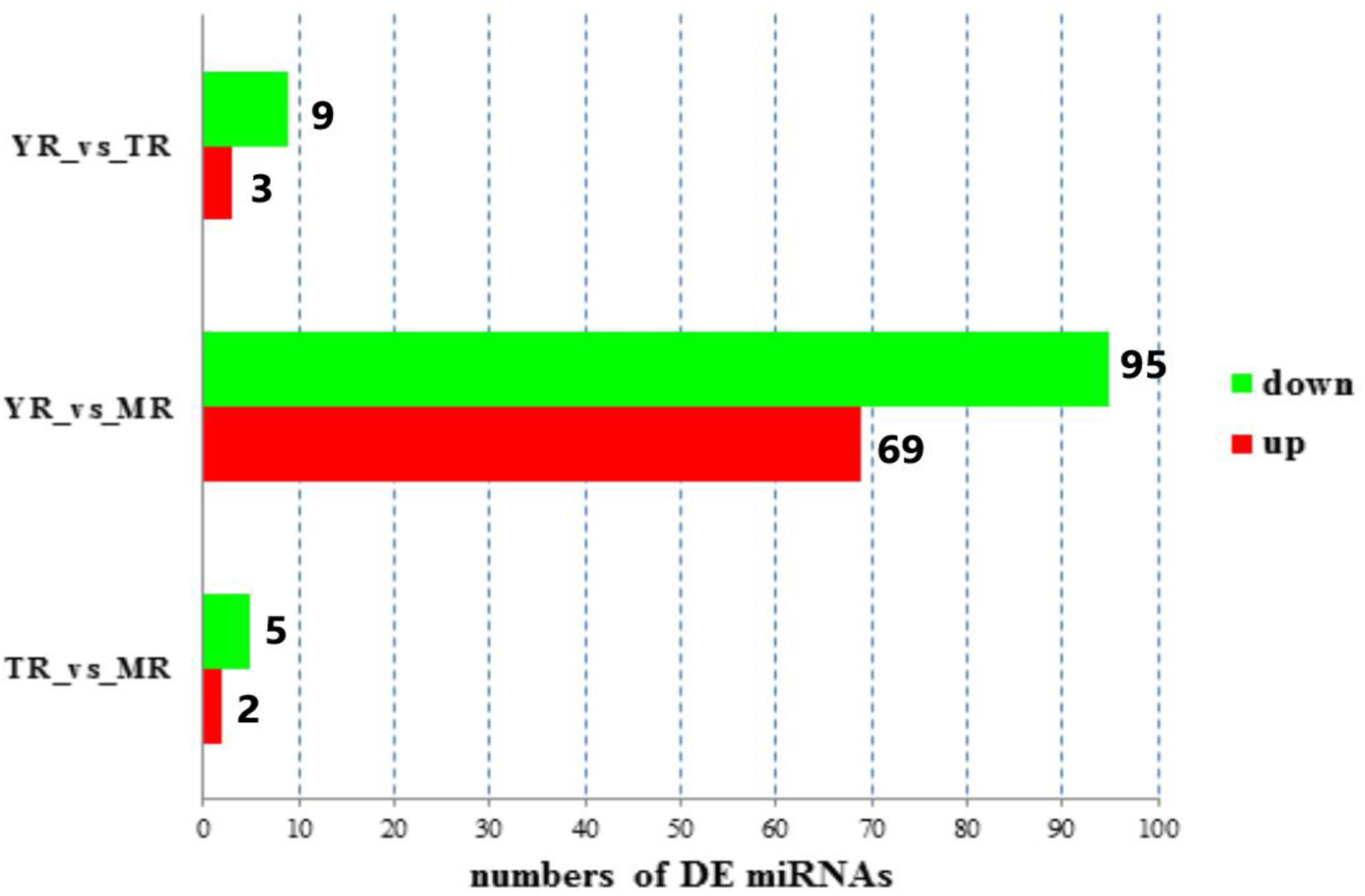
Up and down-regulated miRNAs in the rabbit perirenal adipose during the three growth periods (*P* < 0.05).

To intuitively understand the expression of DE miRNAs in YR-vs-MR, hierarchical clustering was performed on the 164 screened miRNAs (**Figure 3**). As shown in Figure 3, 164 miRNAs showed differential expression patterns according to the different growth stages, and libraries of each group were comparable. The number of highly expressed DE miRNAs (red) in the MR group was significantly higher than the YR group.

**Figure 3.**
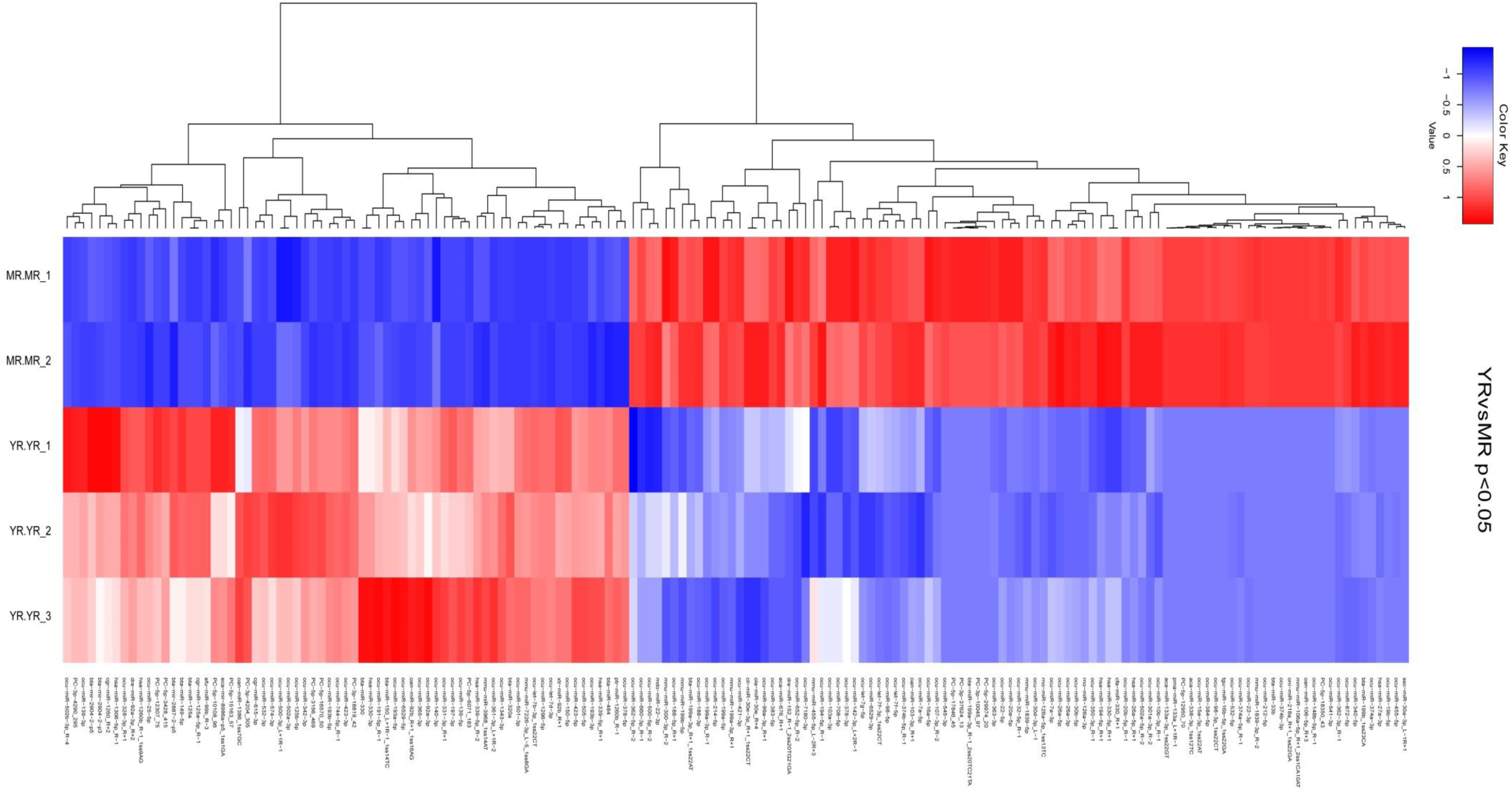
Hierarchical clustering analysis of the miRNA expression profiles from YR-vs-MR with 164 DE miRNAs.

### Enrichment analysis of the target genes of DE miRNAs

Target genes of the DE miRNAs were predicted using TargetScan and miRanda software, and their intersections were taken as final target genes. The number of targets of the 174 DE miRNAs were 13,204. According to the relationship between miRNAs and their target genes, the GO enrichment analysis showed that 13,347 GO terms were obtained, including 8807 terms of biological process (BP), 1279 terms of cell component (CC), and 3261 terms of molecular function (MF), amongst which 1048 terms were significantly enriched (*P*<0.05). Analysis of the 1048 GO terms showed that the target genes of DE miRNAs were significantly enriched in protein binding, cytoplasm, zinc ion binding, regulation of cell growth, and ATP binding (**Figure 4**).

**Figure 4.**
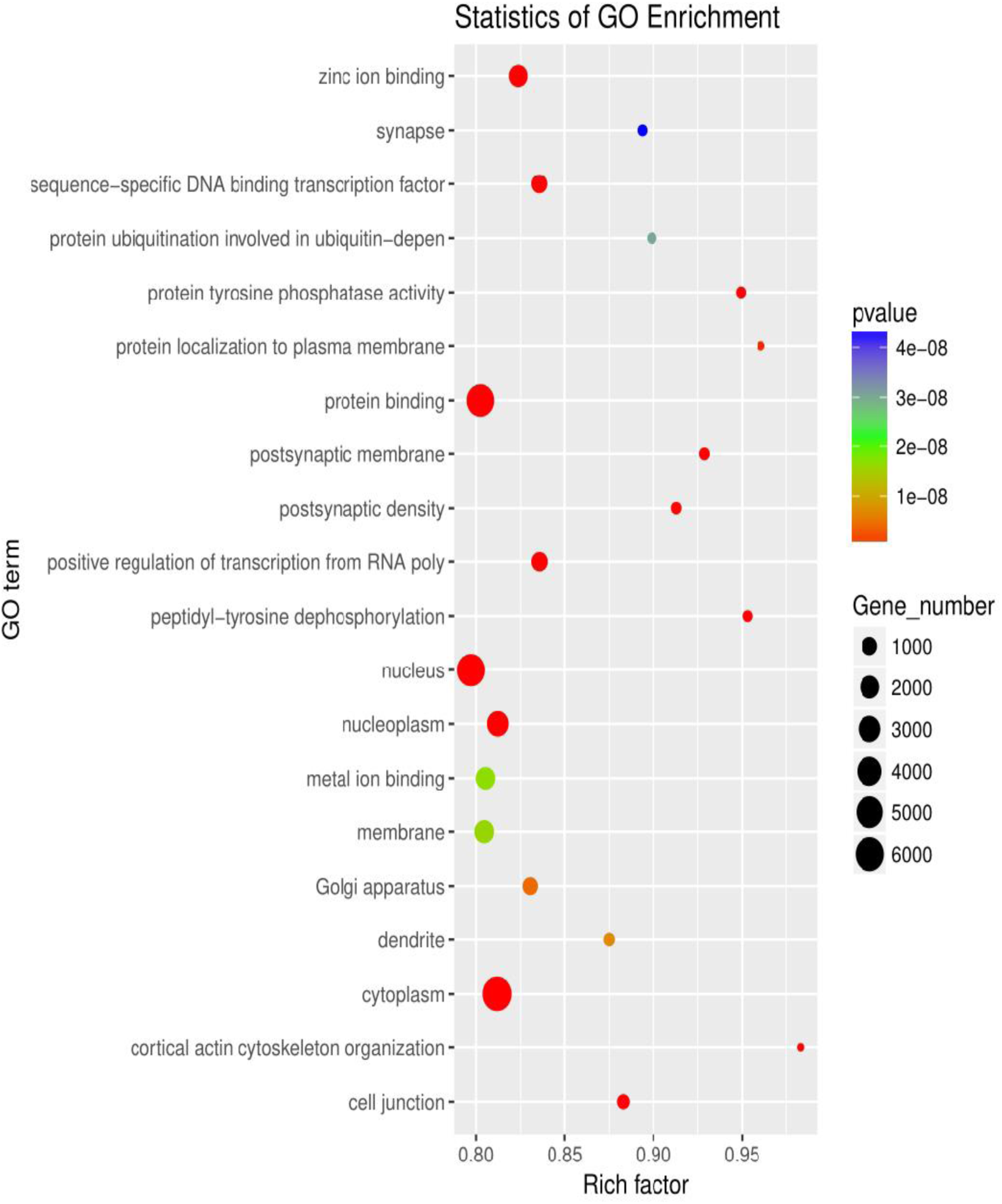
Top 20 significant terms of GO enrichment analysis of target genes of DE miRNAs at *p*-value < 0.05.

To more comprehensively describe the functions of the target genes during the different growth stages, enrichment analysis of the KEGG pathways was used to understand the biological functions of the genes. The results found that the target genes of DE miRNAs were enriched to 315 KEGG pathways, 91 of which were significantly enriched (*P*<0.05), including the MAPK signaling pathway, Wnt signaling pathway, Renin secretion, FoxO signaling pathway, and Aldosterone synthesis and secretion (**Figure 5**).

**Figure 5.**
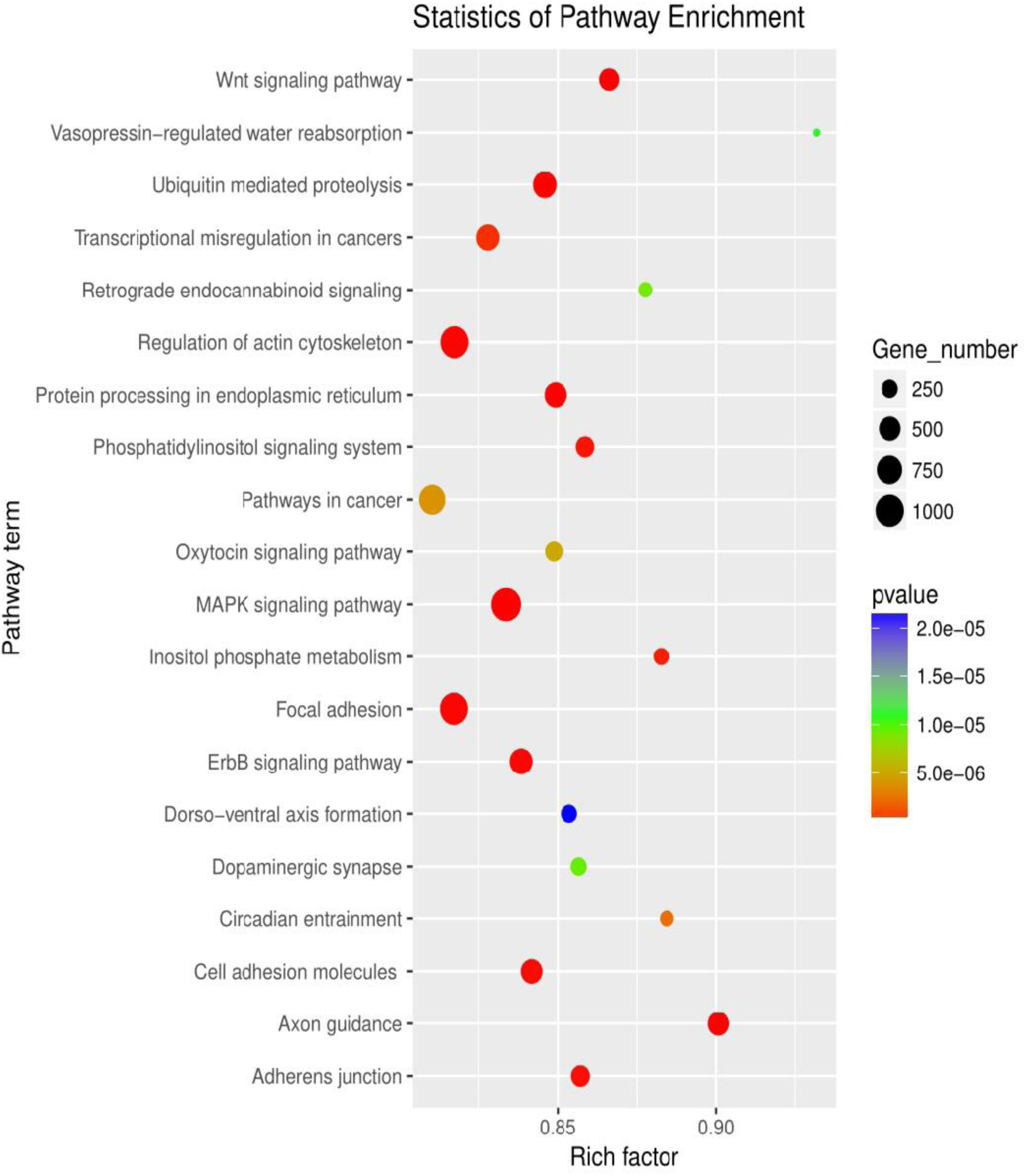
The top 20 significant terms of KEGG Pathway analysis of target genes of DE miRNAs at *p*-value < 0.05.

### Validation of DE miRNAs

To validate the reliability of the miRNA-seq data, six miRNAs (ocu-miR-1296-5p, ocu-miR-193b-3p, mmu-miR-3968_1ss14AT, mmu-miR-199a-3p_R+1, ocu-let-7d-3p, ocu-miR-7a-5p) were randomly selected from 174 DE miRNAs to validate their expression profiles at these three growth stages by q-PCR. The results showed that all six miRNAs were differentially expressed during the different growth stages. In addition, the six miRNAs exhibited a similar trend between the results of miRNA-seq and q-PCR (**Figure 6**). Therefore, the FPKM obtained from the miRNA-seq datasets can be reliably used to determine miRNA expression, and confirmed the importance of DE miRNAs during the growth of rabbit adipose tissue.

**Figure 6.**
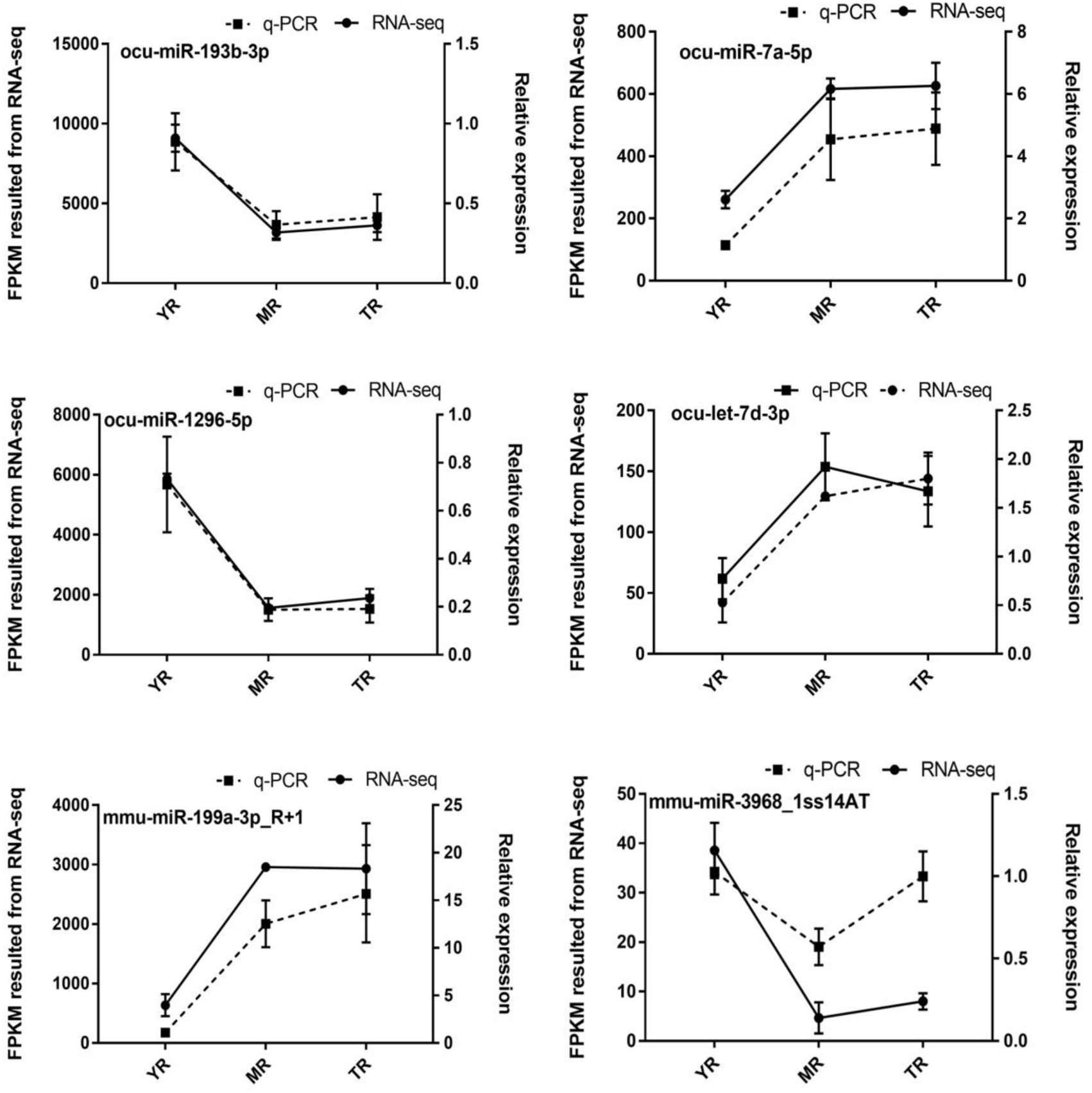
Validation of the six randomly selected DE miRNAs by q-PCR.

## Discussion

In eukaryotes, miRNAs act as a broad class of widely occurring small-molecule ncRNAs that regulate gene expression though targ eting mRNA transcription degradation and translation(Cesar et al. 2002; Xuemei 2004). MiRNAs play important roles in animal growth and development, host immune responses, adipose differentiation and lipid metabolism. Currently, approximately 2,000 miRNAs are recognized in human and mouse genomes, the majority of which are expressed in a tissue-dependent manner(Ana and Sam 2014; Mariana et al. 2002). However, studies on the regulation of miRNAs during rabbit adipose growth and development are lacking. Here, we used MiRNA-sequencing to identify 987 miRNAs during three important stages of rabbit adipose growth, including 780 known miRNAs an d 207 newly predicted miRNAs. The miRNAs were derived from 1 03 families with 643 seed region specificities, including miR-30 and miR-204. Studies have shown(Zaragosi 2011) that miR-30a and mi R-30d induce lipogenesis in obese patients through targeting *RUNX2* and miR-30c respectively, promoting the differentiation of human adipocytes(Karbiener et al. 2011).

We compared the identified miRNAs to other species, which were distributed into 67 miRNAs that included has, mmu, and bta. In-depth analysis of the obtained miRNAs lengths revealed that both known miRNAs and newly predicted miRNAs were mainly 22 nt in length, and increased in abundance during the three growth stages. Similarly, using HiSeq sequencing, Wang and colleagues(Wang et al. 2017) identified 329 known miRNAs and 157 new miRNAs during the development of porcine adipose. Additionally, Wang and coworkers(Wang et al. 2018b) identified 105 DE miRNAs through the deep sequencing of duck adipose tissue and differentiated proadipocytes *in vitro*, demonstrating that miRNA expression varies among different species.

In the present study, the DEGseq R language package was used to identify DE miRNAs. We identified 174 DE miRNAs during the three growth stages of rabbits that were mostly down-regulated. Comparison of each of the stages showed that the number of DE miRNAs at 35 day and 85 day were highest. Adipose growth in the rabbits was significantly affected by age and miRNA expression was more prevalent during early growth stages. Amongst the 174 DE miRNAs, some were distributed in miR-133, miR-30 and let-7 families. Related studies showed that miR-133a is expressed in brown and white adipose tissue, directly targeting the 3’UTR region of *Prdm16*(Weiyi et al. 2013). miR-let-7b regulates the levels of human adipose tissue-derived mesenchymal stem cells (hAT-MSCs), and the transient inhibition of miR-let-7b enhances the differentiation of hAT-MSCs(Effat et al. 2015). These results suggest that the DE miRNAs identified in this study play regulatory roles during adipose growth in rabbit.

MiRNAs pair with the 3’UTRs of target genes to inhibit translation and silence gene expression at the post-transcriptional level. Bioinformatics estimates that 30-80% of the mammalian miRNAs target multiple cellular mRNAs(Friedman et al. 2008). In general, target genes regulated by the same miRNA originate from the same gene family(Yang et al. 2013). In this study, the 174 DE miRNAs were predicted to target 13,204 genes, with an average of 76 genes targets for each predicted miRNA. Moreover, the target genes regulated by single miRNAs originated from the same family, and the DE miRNAs showed obvious temporal characteristics.

Compared to lncRNAs(Wang et al. 2018a), miRNAs and the target genes of DE miRNAs were mainly involved in GO functional terms including metabolic process, cell process and single organism process in the classification of biological processes, partial cells and organisms in the classification of cell components, and binding and catalytic activity in the classification of molecular functions. Based on our in-depth analysis of the 1048 significantly enriched GO terms, it was found that amongst the top 10 GO terms of biological process, cell composition and molecular function, some terms that strongly promote growth and volume increases in adipocytes, including protein localization to the plasma membrane, protein ubiquitination involved in ubiquitin-dependent protein catabolic process, regulation of cell growth, cytoplasm, Golgi apparatus, membrane, protein binding, protein tyrosine phosphatase, zinc ion binding, ATP binding, and cadherin binding were identified. However, there were few related terms regarding glyceric acid absorption and lipid droplet formation during adipose hypertrophy. Recent studies on miRNA expression in human adipose tissue found that the expression of miRNAs were specific to the site of adipose tissue(Nora et al. 2009; Ortega et al. 2010). Some miRNAs were associated with adipose tissue morphology, adipocyte size, and metabolic functions (fasting glucose, triglyceride). Combined with our data, the target genes of DE miRNAs more highly influenced cell membrane growth, protein synthesis and utilization, energy utilization and transformation, but their role in lipid droplet accumulation in adipocytes was not obvious.

Amongst the 91 pathways significantly enriched by KEGG, MAPK signaling pathway, Wnt signaling pathway and aldosterone synthesis and secretion pathways have been shown to regulate the growth and development of adipocytes. Related studies have shown that some miRNAs participate in the regulation of adipose deposition, for example, miR-148a promotes adipose synthesis by inhibiting the expression of *Wnt1*(Shi et al. 2015), whilst the over-expression of miR-10b L20 significantly increases the levels of adipose and triglyceride(Lin et al. 2010). In addition, miRNA families such as let-7, miR-30, miR-17, miR-148(Jing et al. 2012) and miR-24(Qiang et al. 2008; Kang et al. 2013) are involved in animal adipose deposition. miR-20a regulates adipocyte differentiation by targeting lysine-specific demethylase 6 b and transforming growth cytokine β signal(Zhou et al.). Previous studies(Michael et al. 2009) assessed the anti-adipogenesis characteristics of miR-27b, which was down-regulated during adipocyte differentiation and weakened the induction of *PPARc*. The expression of miR-95 significantly correlated with adipocyte size, and its expression significantly increased during adipocyte differentiation(Nora et al. 2009). Therefore, the results of this study suggest that miRNAs with tissue and developmental stage specificity play key roles in the growth and maturation of rabbit adipose tissue.

## Conclusions

In conclusion, to the best of our knowledge, this is the first report to perform miRNA profiling of rabbit perirenal adipose tissue during different growth stages, which identified 987 miRNAs and 174 DE miRNAs associated with adipogenetic pathways. These included the regulation of cell growth, zinc ion binding, MAPK signaling pathway, and Wnt signaling pathway. These DE miRNAs therefore regulate the growth and hypertrophy of adipose tissue in rabbits.

## Acknowledgments

We thank the staff at our laboratory for their ongoing assistance. We also thank Xing-zhou Tian for insightful feedback on the study.

## Authors’ contributions

GZW, XBJ, SJL designed and directed the study. GZW, GG, KD performed the experiments, data analysis and drafted the manuscript. XTT, JXM, SYC contributed to the analysis and writing of the manuscript. SQH, JW, SJL critically reviewed drafts of the manuscript and made comments to improve clarity. All authors approved the final version of this article.

## Animal ethical approval

All surgical procedures involving rabbits were performed according to the approved protocols of the Biological Studies Animal Care and Use Committee, Sichuan Province, China. Rabbits had free access to food and water under normal conditions and were humanely sacrificed as necessary to ameliorate suffering.

